# Graphite: painting genomes using a colored De Bruijn graph

**DOI:** 10.1101/2023.10.08.561343

**Authors:** Rick Beeloo, Aldert L. Zomer, Bas E. Dutilh

**Affiliations:** Theoretical Biology and Bioinformatics, Utrecht University, Padualaan 8, 3584 CH Utrecht, the Netherlands; Department of Infectious Diseases and Immunology, Faculty of Veterinary Medicine, Utrecht University, 3584 Utrecht, the Netherlands; Institute of Biodiversity, Faculty of Biological Sciences, Cluster of Excellence Balance of the Microverse, Friedrich Schiller University Jena, 07743 Jena, Germany

**Keywords:** chromosome painting, *Campylobacter*, antibiotic resistance genes, horizontal gene transfer, bioinformatic tool, suffix array

## Abstract

The recent growth of microbial sequence data allows comparisons at unprecedented scales, enabling tracking of strains, mobile genetic elements, or genes. Querying a genome against a large reference database can easily yield thousands of matches that are tedious to interpret and pose computational challenges. We developed Graphite that uses a colored De Bruijn graph (cDBG) to paint query genomes, selecting the local best matches along the full query length. By focusing on the closest genomic match of each query region, Graphite reduces the number of matches while providing promising leads for genomic forensics. When applied to hundreds of *Campylobacter* genomes we found extensive gene sharing, including a previously undetected *C. coli* plasmid that matched a *C. jejuni* chromosome. Together, genome painting using cDBGs as enabled by Graphite, can reveal new biological phenomena by mitigating computational hurdles. Graphite is implemented in Julia, available at https://github.com/MGXlab/Graphite.

## Introduction

Tracking genetic sequences is crucial for understanding various biological phenomena, such as recombination and horizontal gene transfer (HGT) of genetic elements, as well as source attribution in epidemiology. Advances in DNA sequencing, coupled with the abundance of sequencing data in databases, enable us to reconstruct the movement of genetic information at different scales. For example, the rearrangement of chromosomes in a single cell^1^, exchange of plasmids or conjugative elements between different bacteria^2,3^, movement of plasmids through multiple environments^4,5^, and following microbes across continents^6,7^. Each genome or metagenome sequence serves as a snapshot at a specific time and place. Comparing these snapshots allows us to identify closely related sequences across geographic and ecological boundaries, and track genetic information at different scales, from within a patient to across the globe.

Several mechanisms can lead to the repeated observation of a given microbial sequence. First, the observed strains may be very closely related, suggesting a recent physical exchange between the sampling sites, or both from a third source^8–10^. Second, the sequence may reflect a conserved mobile genetic element (MGE), which is transferred between different microbial lineages independently of vertical inheritance^11^. Either way, the timescale of the events may be inferred from the similarity between the two sequences. Just after such events the sequences should be highly similar, but over time they increasingly accumulate mutations such as point mutations, loss of segments, and insertion of new information via recombination^11,12^. From the perspective of a given query, longer sequence matches reflect more recent exchanges. Thus, searches for the longest stretch of sequence identity in a database of target sequences can be used to identify the closest genomic link of the query.

Given the expanse and rates of exchange of DNA, especially in prokaryotes^13,14^, there can be different matches across a single chromosome. To establish such links, chromosome painting has been used within cells and across species. Originally, chromosome painting was performed experimentally, by probing specific regions of the chromosome with fluorescently labeled DNA^15^, and the resulting color patterns were used to compare chromosome structures. In case of genomic exchange or chromosomal rearrangements, the sequence segments can be traced back to their source based on their color. Experimental chromosomal painting has been applied to a variety of species such as plants^16,17^ and fungi ^18^.

*In silico* chromosome painting similarly identifies genomic links between a query sequence and reference database, and has been used to study the population structure in, e.g., humans^19^ and *Helicobacter pylori* ^*20*^. From a genomic forensics perspective, identifying the longest identical match of a given query segment would yield the closest genomic link, within the search frame of a given reference database. For example, in the case of a recent geographic migration or HGT, the best match might reflect the most likely origin or donor of a subsequence. Identical matches between two sequences are commonly defined as a maximal exact match (MEM), which cannot be extended without introducing mismatches^21^. Algorithms to identify MEMs can broadly be categorized into those using suffix arrays and their derivatives^21–24^, and those that use hashing^25,26^. MEM identification usually starts with partitioning the reference sequences into k-mers, which are indexed and queried for k-mers of the query sequence. Both methods, especially suffix arrays, are memory intensive when applied to large reference datasets. This was circumvented by bfMEM by only indexing k-mers present in the query^26^. These tools have shown excellent performance in resolving pairwise MEMs between a query and a large reference dataset. Still, pairwise comparisons of thousands of genomes remain computationally infeasible.

One data structure allowing efficient large-scale sequence comparison is the colored de Bruijn graph (cDBG). To construct a cDBG all the sequences are divided into k-mers that are used to build a de Bruijn graph, where edges represent the overlaps between adjacent k-mers in the sequences. The optimal k-mer size greatly depends on the question but generally ranges from 18-31 nucleotides^27^. Each sequence used to build the cDBG traverses a specific path of nodes that overlaps with other paths in regions of sequence identity. To further compress the cDBG, non-branching paths are merged into unitigs, resulting in a compacted cDBG (ccDBG) whose nodes are equal to or bigger than the original k-mer size. The compression feature of ccDBGs, combined with their ability to enable fast sequence comparisons, makes them suitable for tasks including sequence alignments^28^, variant calling^29,30^, homology detection^31^, and genome-wide association studies^32^. State-of-the-art ccDBG construction tools such as CuttleFish can process enormous amounts of sequences^33^. Most graph builders output a Graphical Fragment Assembly format (GFA), which represents the graph and serves as a widely accepted input for tools utilizing ccDBGs. This promotes efficient data sharing by removing the need to build separate indexes for every tool, which is often the most time-consuming part^34^.

Encoding a set of sequences into a ccDBG also implicitly resolves shared regions between the input sequences. For example, if two input sequences jointly traverse the same consecutive set of nodes, this indicates a shared identical subsequence or MEM. To identify MEMs for any given query, its path can be checked for intersections with paths of other sequences. In a large database, one query region could yield MEMs with thousands of different reference sequences, while we are often only interested in the longest match overall. Here we developed Graphite to find the longest MEM (LMEM) for each region in the query using a CDG. Resolving these LMEMS locally across a query provides unique insights into the origin of genetic segments that remain hidden in the wealth of matches. We demonstrate this by applying Graphite to a collection of *Campylobacter* genomes where we zoom in on two LMEMs that suggest an inter-species transfer of genetic material between *C. coli* and *C. jejuni* and *C. coli* and the emerging pathogen *C. hyointestinalis subsp. hyointestinalis*^*35*^

## Results

We developed Graphite to identify the longest maximum exact matches (LMEMs) between one or more queries and multiple reference sequences. Below, we first validate LMEM identification by comparing our results to MEMs identified by E-MEM. Then we use an example dataset composed of 576 *Campylobacter* genomes to show how LMEMs can drastically reduce the number of matches and thus the output file size, and demonstrate how they shed light on relevant evolutionary events.

### Validation of LMEM identification on three bacterial genomes

Graphite uses a ccDBG as input, in this case, we built them using Cuttlefish with a k-mer size of 31 nucleotides^33^. Node sizes in the ccDBG depend on the sequence variation in the input sequences. Mutations such as SNPs introduce bubbles in the cDBG and prevent merging adjacent k-mers into longer sequences during graph compaction. As nodes may be shared by all sequences that overlap by at least 31 nucleotides, a shared node rarely forms the entire MEM, as variations often occur in sequences other than those being compared (**Figure 1a**). Graphite locates the shared nodes between a query sequence and each reference to bidirectionally extend these resolving a MEM. As all references in the database are compared, a given query region will match multiple MEMs, of which Graphite only retains the longest. To validate LMEM selection by Graphite we used both E-MEM and Graphite to query *C. jejuni* CP071576 against the CP071584 and CP085965 genomes. E-MEM identified 1,586 MEMs with CP071584, and 10,990 MEMs with CP085965. Graphite resolved 1,564 LMEMs, most selected from CP071584 with some exceptions (**Figure 3b**). **Figure 3c** shows how these exceptions support Graphite’s LMEM selection algorithm.

**Figure 1.**
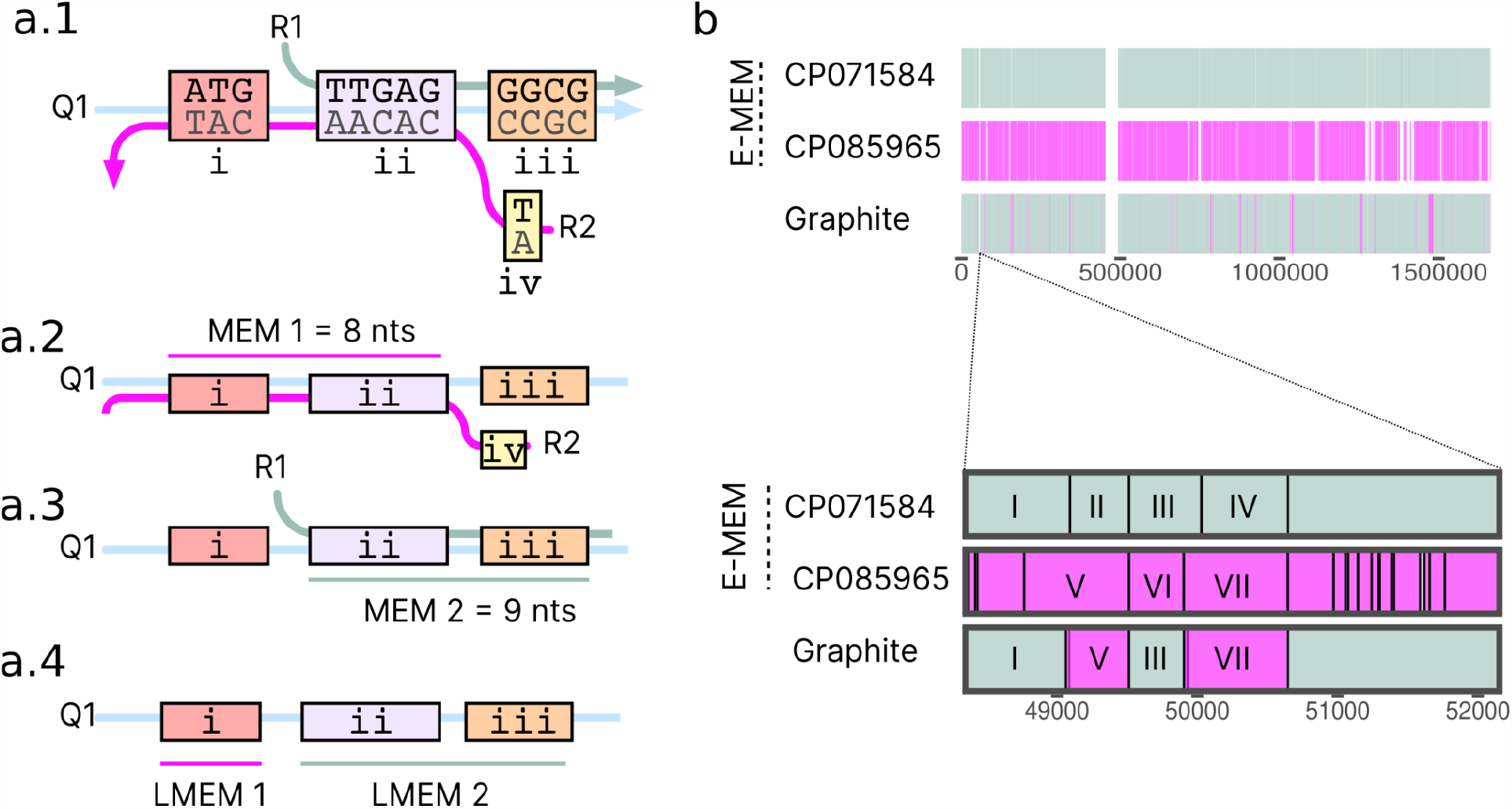
Longest maximum exact match (LMEM) selection. (A) In a compacted graph, the variation in the overall sequence set determines the sequence fragmentation and node size. **a**.**1** shows a toy graph of three sequences Q1, R1, and R2 that overlap in four nodes i-iv (see **Figure S1** for further details). The k-1 overlap between nodes as seen in ccDBGs is not shown for simplicity. The sequence strands are indicated with arrows, the forward and reverse sequences are shown in the nodes. While Q1 and R2 share a MEM of 8nt (**a**.**2**) consisting of nodes i-ii, it does not show up as a single node since another sequence, R1 contains a subsequence of this MEM (node ii, **a**.**1**). Similarly, Q1 and R1 share a MEM of 9nt (**a**.**3**) consisting of nodes ii-iii. **a**.**4** shows how two LMEMs are resolved for this example. Each node is assigned to the longest MEM it is part of. Node ii is part of two MEMs and gets merged with node iii into LMEM 2 (9nt) which is 1 nucleotide longer than MEM 1 (8nt). As the remaining node i is only covered by MEM 1, it is assigned to LMEM 1, and the query sequence is painted accordingly (**a**.**4**). (**b**) To validate Graphite on real data we aligned *C. jejuni* CP071576 against CP071584 and CP085965 using E-MEM and Graphite. The majority of Graphite LMEMs originated from CP071584 (gray). (C) Close-up example of LMEM selection. First, MEM I is selected from CP071584 as it is longer than the multiple overlapping CP085965 MEMs. A part of MEM V is selected as the next LMEM, as V is longer than II. Likewise, a part of MEM III is selected over MEM VI before Graphite’s LMEMs continue into VII.

### Graphite efficiently identifies LMEMs in hundreds of genomes

To test Graphite on a medium-sized real world dataset, we downloaded all 579 complete *Campylobacter* genomes from BV-BRC^36^. After curation 576 remained (see Online Methods, **Figure S3, Table S1**) which were used to construct a ccDBG using Cuttlefish^33^ with 3,496,408 nodes and 115,691,195 edges. The genomes spanned 33 different species (excluding *Campylobacter* sp.) with the majority of genomes being derived from *C. jejuni* (n=363, 63%) and *C. coli* (n=86, 15%), while 29 species were represented by less than ten genomes. Next, we compared the performance of different tools for finding MEMs including E-MEM^25^, Bf-MEM^26^, and LMEMs by Graphite, querying all 86 *C. coli* genomes against the remaining 490 *Campylobacter* genomes **(Table 1**). Graphite ran almost four times faster than the next tool, E-MEM. Graphites runtime includes the time needed to construct the ccDBG (∼1.5 minutes), but unlike the MEM finders, the ccDBG can be reused for additional queries and other analyses such as variant calling and visualization. By focusing on finding LMEMs rather than all MEMs, we reduced the output by over 50x (**Table 1**, matches ≥31nt by Graphite and E-MEM). This demonstrates that LMEMs can significantly save storage space, especially when querying big reference datasets. As we are often only interested in the longest matches, for example for *in silico* chromosomal painting or HGT analysis, Graphite obsoletes post-processing of vast numbers of MEM results by already selecting the longest MEM during alignment.

**Table 1.**
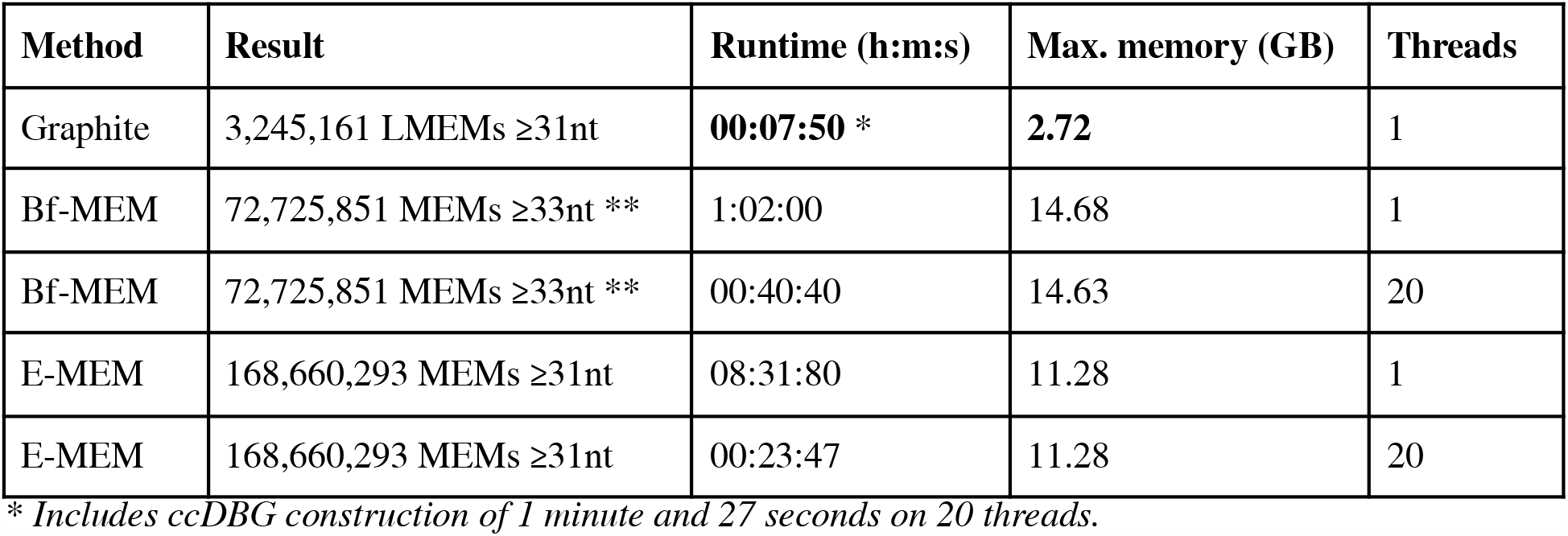
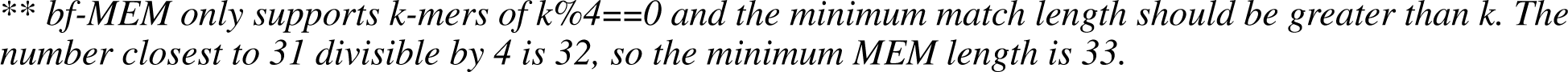
Performance comparison of graphite and two MEM finding algorithms when querying 86 *C. coli* versus 490 other *Campylobacter* genomes using Graphite and two MEM finding algorithms.

### Discovery of unique nodes in the Graphite ccDBG reveals unexpected genomic entities

Generally, short nodes in the ccDBG were shared by more genomes than long ones (**Figure 2**). For example, nodes present in at least half of the genomes (>=288) were on average 64 nucleotides (nt) long with the longest being only 112nt. Of all 3.5 million nodes, 24.8%, 9.6%, and 0.07% were present in at least 10, 100, and 500 genomes, respectively. The abundant sharing of the nodes between different genomes illustrates how nodes can function as anchor points for MEM and LMEM detection. The top five longest nodes in the graph exceeded 25,000nt and were all present in a single genome. These large nodes represent non-branching paths in the graph, i.e. consisting of unique 31-mers that are never found in another genome in the dataset. They originated from *C. jejuni* (42,403nt), *Campylobacter sp*. (31,087nt) as well as under-represented species *C. curvus* (29,437nt, four genomes), and *C. concisus* (28,388nt and 27,886, eight genomes). The unique *C. jejuni* node corresponded to a sequence annotated as plasmid pFORC_083_2, found in *C. jejuni* strain FORC_083 that was isolated from chicken in South Korea. It is interesting to note that this plasmid probably represents a bacteriophage, as it encodes typical bacteriophage genes including structural and replication clusters, integrase, and a terminase large subunit that is related to sequences found in various *Firmicutes* (**Figure S4**), while *Campylobacter* belongs to the *Epsilonproteobacteria* class. It is also predicted as a member of the *Caudoviricetes* class by IMG/VR ^37^ and was found to match a 30nt CRISRPR spacer in the firmicute *Lactobacillus delbrueckii* subsp. *delbrueckii* strain KCTC 13731 with three mismatches (90% identity) by CRISPRimmunity^38^. Whether this is a case of contamination in this finished genome sequence or a bacteriophage with an unusually wide host range remains to be seen^39^, but the example illustrates that the detection of long, unique nodes in a pan-genome graph may quickly reveal unexpected or foreign sequences for follow-up.

**Figure 2.**
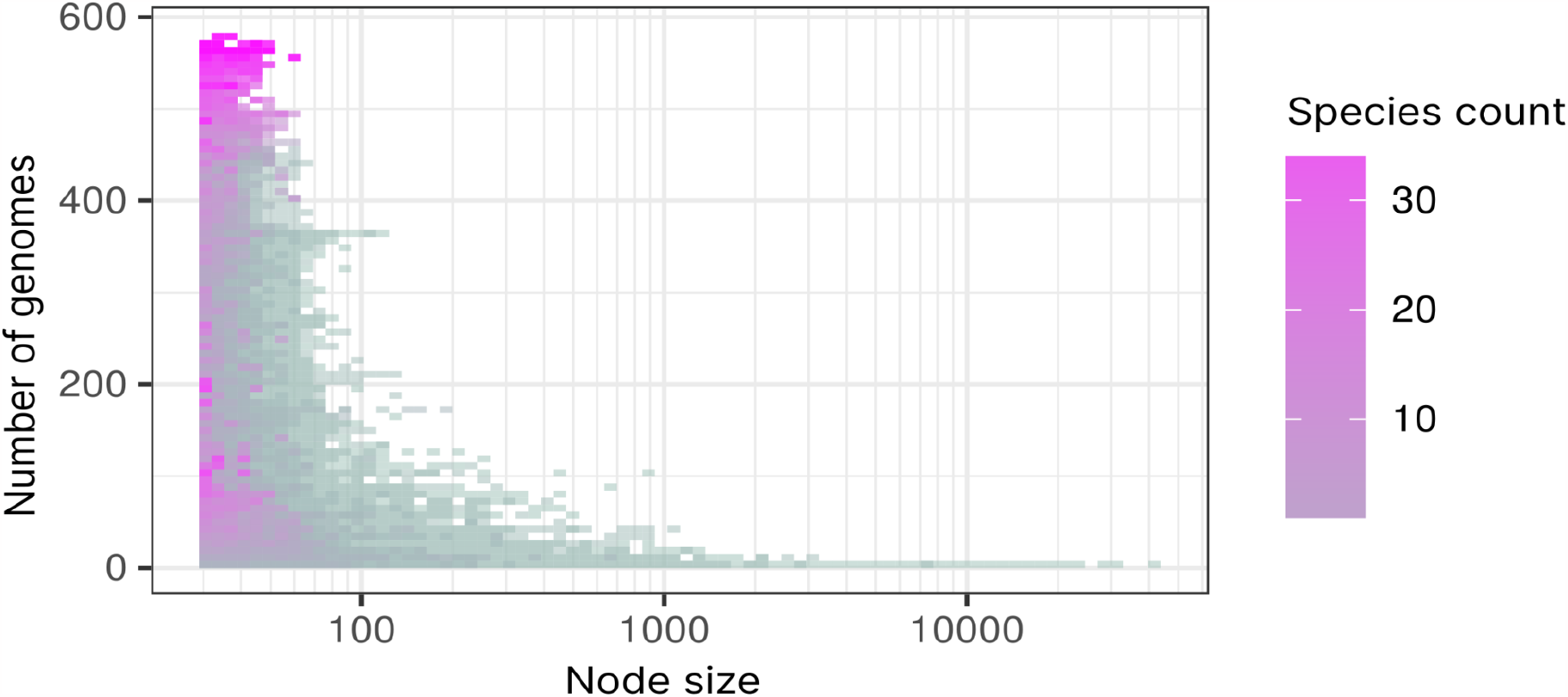
Relation between the size of nodes and the number of genomes and species they occur in. Short nodes may occur in many genomes and species, while long nodes over 120nt rarely occur in more than 100 genomes.

### LMEMs quanitfy interspecies genetic transfer within the *Campylobacter* genus

*Campylobacter* is known to actively engage in horizontal gene transfer (HGT) via various mechanisms, such as the uptake of environmental DNA and the exchange of plasmids^40^. To demonstrate how LMEMs can contribute to the detection of such genomic links and zoom in on long regions that are likely recently acquired, we queried each individual genome against all others using Graphite, which took 5 hours and resulted in 2,109,030 LMEMs. One advantage of LMEMs is that they form a one-dimensional output containing, for each query region, the longest match in the database. This can be plotted to obtain a genome painting visualization for every genome in the *Campylobacter* genus, which reveals substantial HGT (**Figure 3**). When considering all LMEMs, 5.29% of the *C. coli* sequence had the closest match in *C. jejuni*, and 2.64% of the *C. jejuni* DNA had the closest match in *C. coli*. The frequencies of shared DNA between these two species varied greatly depending on the contig, ranging from 0 to 99.93% for *C. jejuni* and from 0.000112% to 99.81% for *C. coli*. While high frequencies were mainly observed for shorter contigs (**Figure 3, Figure S5**), the *C. coli* chromosome CP092025 matched 19% of its DNA with *C. jejuni* and the *C. jejuni* chromosome CP059964 matched 10% of its DNA with *C. coli*. These results confirm the high transfer frequency of genetic material between *C. coli* and *C. jejuni*.

**Figure 3.**
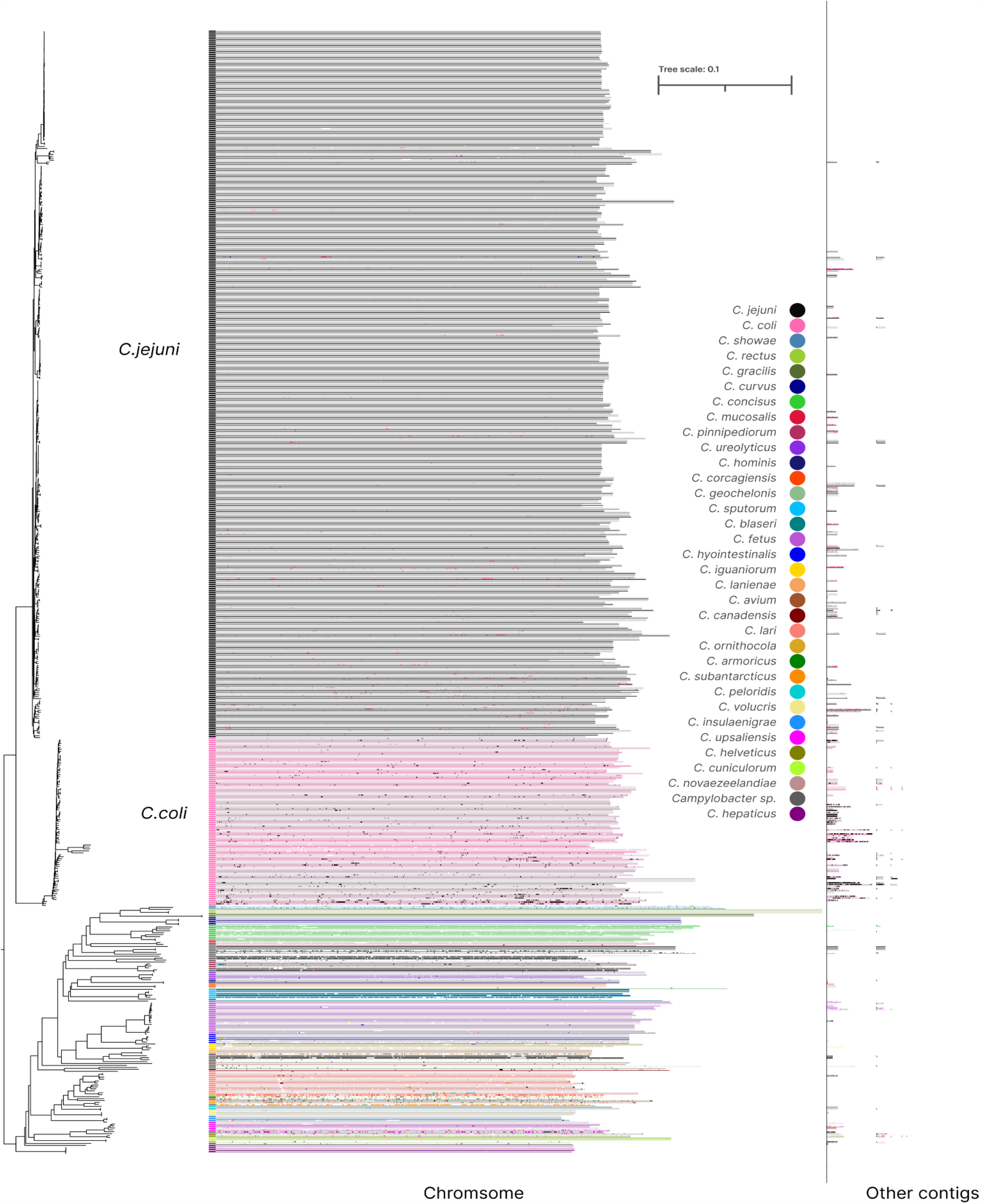
LMEM painting of 576 Campylobacter genomes. On the left is the Masthree clustering (**Figure S3**). To the right, each row represents a query genome along a colored line representing the species it belongs to. Based on the LMEMs, the genomes are painted above the line, colored according to the matched species, or gray when the LMEM was found in the same species. For example, *C. coli* LMEMs matching *C. jejuni* are black in the *C. coli* clade, while *C. jejuni* LMEMs matching *C. coli* are pink in the *C. jejuni* clade.

### LMEMs highlight biologically relevant genomic transfer events

To further explore our results, we investigated long LMEMs that were shared by genomes from different species in search of striking cases of HGT. One LMEM of 16,417nt was shared between a *C. jejuni* chromosome (CP107256) and a *C. coli* plasmid (CP013035, **Figure 4a**). This LMEM encoded several proteins including the *tetO* gene conferring resistance against tetracycline antibiotics. Inter-species LMEMs were also observed among other species (**Figure 3**), for example, we detected a 2,068nt LMEM between *C. jejuni* (CP047082) and *C. hyointestinalis subsp. hyointestinalis* (CP015575), which encodes an ISChh1 transposon according to ISfinder ^41^ that disrupted the *pycB* gene in *C. hyointestinalis subsp. hyointestinalis* (**Figure 4b)**. We also explored other paths in the cDBG that traversed the LMEM nodes, showing that the transposon was present in twenty genomes in a variety of different contexts (**Figure 4b**).

**Figure 4.**
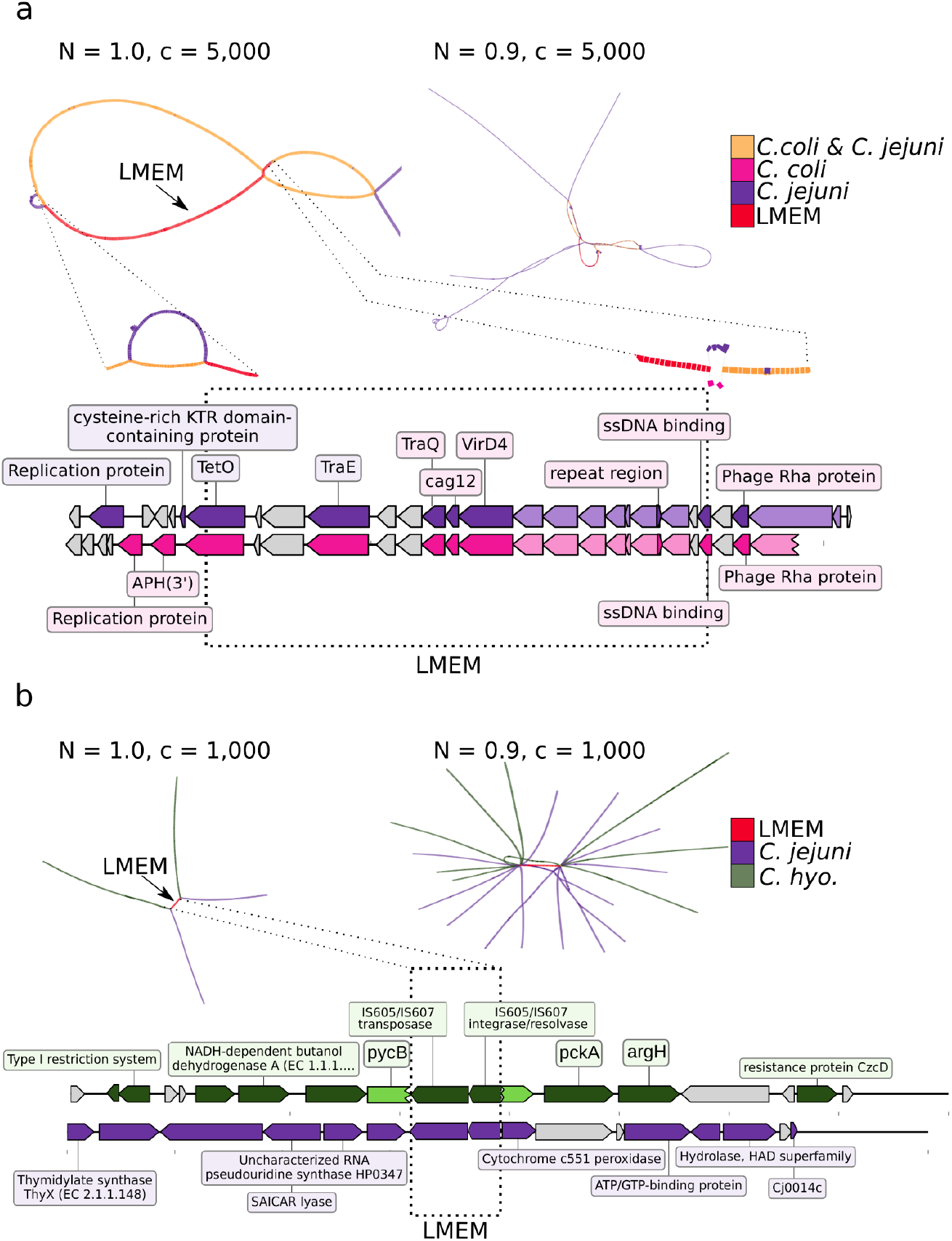
Example subgraphs of two LMEMs. (**a**) A 16,417 nt LMEM between a *C. coli* plasmid (CP013035, positions 23,541-39,957) and *C. jejuni* chromosome (CP107256:1,208,016-1,224,432). The parameter c specifies the fraction of LMEM nodes that a path has to traverse in order to be included in the subgraph visualization and n reflects the number of flanking nodes to display (see online methods). The left panel in A shows the subgraph of the LMEM (n=1.0) with two genomes, 11,045 nodes, and 11,057 edges. The right panel shows a relaxed subgraph of the LMEM with paths sharing ≥90% (n=0.9) of the LMEM nodes. This relaxed subgraph contained 23,051 nodes and 23,665 edges, with paths from 18 genomes. In the left panel we zoom in on the bubbles that prevent further LMEM extension and for example, see that *C. coli* had aminoglycoside resistance gene (APH(3’)) flanking the LMEM which was absent in *C. jejuni* at this position. The gray-colored genes were annotated as hypothetical by BV-BRC and the light-colored genes encoded VirB proteins but their annotations were omitted for clearer visualization. (**b**) A subgraph with 4,103 nodes and 4,102 edges for a 2068nt LMEM encoding a transposon was found in two genomes from *C. jejuni* (CP047082: 58,822-60,889) and *C. hyointestinalis subsp. hyointestinalis* (CP015575: 1,258,714-1,260,781) abbreviated as *C. hyo*.. For the latter species, the transposon disrupted the *pycB* gene encoding a pyruvate carboxylase according to the NCBI annotation (light green). When we slightly relaxed the subgraph threshold to n=0.9 we obtained a subgraph with 20,255 nodes and 20,270 edges, with paths from 20 genomes. The extensive branching of paths outside the LMEM indicates the diverse genomic context of this transposon in different *C. jejuni* and *C. hyointestinalis subsp. hyointestinalis* genomes.

**Figure 5.**
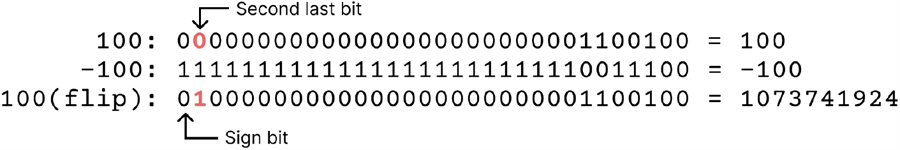
Node bit flipping. Node identifiers in the ccDBG can be either positive or negative, indicating traversal direction. To standardize them as positive, we use the second-to-last bit. For example, node “100” can be traversed forward (100) or in reverse complement (-100). To make -100 positive we can first make it positive and flip the second last bit (0 → 1), compare 100 versus 100(flip)). While 100(flip) binaries code now represents a different number (1073741924) we can derive the original (100) by flipping back the second last bit. This becomes useful when aligning forward and reverse complement matches as the forward and reverse traversal now have the same binary code, except for that second last bit.

**Figure 6.**
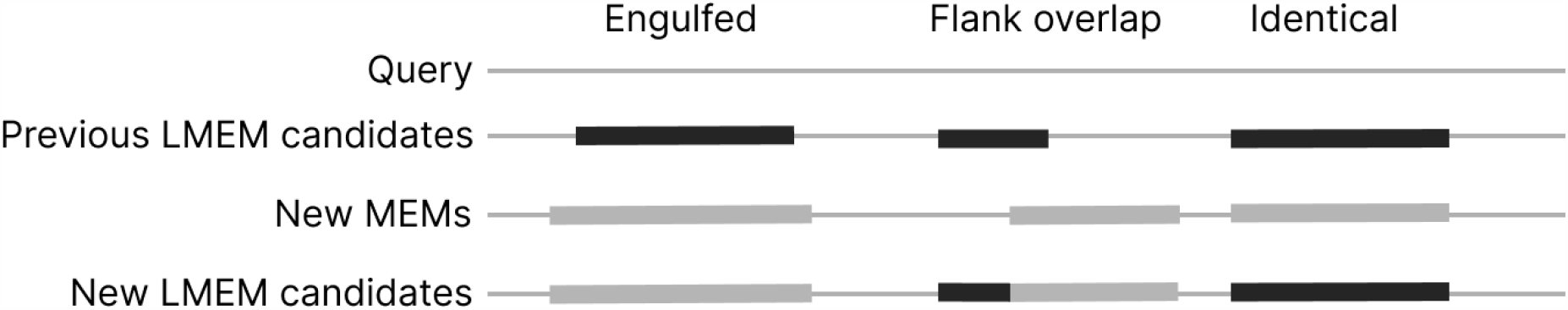
LMEM replacement criteria. Upon discovering a new MEM, we compare its length to the previous LMEM candidates covering the same sites. Three columns indicate different possibilities. If the new MEM is longer or if there is no previous LMEM at those sites, it replaces the current LMEM (engufled). When the MEM and LMEM have overlapping regions, the overlapping region gets assigned to the longest of the two (flank overlap). If a MEM matches the LMEM’s length precisely, we retain the initially found LMEM (identical).

## Discussion

*In silico* chromosome painting allows source tracking of individual query sequences in high throughput. Here we present Graphite, an algorithm that efficiently identifies LMEMs between query and reference genome sequences by exploiting the ccDBG data format. Graphite addresses the challenge posed by traditional MEM finders, which often generate an overwhelming number of MEMs^25^, complicating downstream analysis. For example, while filtering MEMs based on a minimum length can effectively reduce the number of matches, it could also exclude MEMs in regions that are inherently more challenging to align. Graphite overcomes this by locally selecting the LMEM, reducing the number of MEMs by over 50 times compared to two established MEM finders (**Table 1**) while also retaining optimal query coverage (**Figure 1**). Moreover, compared to E-MEM and bf-MEM, Graphite was faster and used less memory (**Table 1**).

We showed that LMEMs not only serve as an output filtering approach but also provide new insights to answer biological questions. For example, by using Graphite to identify LMEMs in an all-versus-all comparison involving 576 *Campylobacter* genomes, we quantified overall inter-species genomic transfer rates and highlighted interesting cases. Notably, inter-species LMEMs were particularly prevalent among shorter contigs (on the right in **Figure 3**), which likely represent plasmids or other mobile genetic elements (MGEs) that are extensively shared within the *Campylobacter* genus^40^. We also found that inter-species LMEMs between *C. coli* and *C. jejuni* were common for chromosomal contigs sometimes covering up to 19% of the sequence. Interestingly, our LMEM analysis also reveals that *C. coli* shared a greater portion of its DNA with *C. jejuni* than vice versa, which is in line with previous observations that suggest a directionality in the transfer between these species^42^. *C. jejuni* was overrepresented in our dataset which might have affected this observation. The abundant exchange of genetic material between *C. coli* and *C. jejuni* has been proposed to result in a process of introgression or despeciation^42^.

Zooming in on the *Campylobacter* results supports the notion that LMEMs can effectively identify inter-species transfers. We found a 16,417nt tetracycline resistance encoding LMEM between a 44,064nt *C. coli* plasmid and *C. jejuni* chromosome (**Figure 4a**). In addition, our subgraph visualization of the LMEM region showed that almost all nodes of the *C. coli* plasmid were present in the *C. jejuni* chromosome. Previously, this particular *C. jejuni* chromosomal segment had been theorized to arise from a conjugation event, wherein a *tetO* gene and a prophage were introduced into a plasmid, possibly from a common ancestor^43^. The fact that Graphite linked this *C. jejuni* region as an LMEM to a *C. coli* plasmid suggests that *C. coli* played a role in this event as a recent donor or recipient of the sequence from *C. jejuni* (**Figure 4b**). Another example is the ISChh1 transposon-encoding LMEM between *C. jejuni* and *C. hyointestinalis subsp. hyointestinalis* (**Figure 4b**). While horizontal gene transfer between these two species is not extensively researched, observations suggest that they engage in recombination when cohabiting in the same environment^44^. Although we did not observe it in this instance, transposons, including ISChh1-like transposons^45^, mediate the transfer of a variety of genes enhancing the pathogenicity of the host, such as resistance genes and virulence genes^46^. In this case, for *C. hyointestinalis subsp. hyointestinalis*, the transposon disrupts the *pycB* gene, potentially pseudogenizing it^47^. When visualizing the subgraph of the transposon, with all paths traversing ≥90% of the LMEM nodes, we found this transposon in twenty different genomes. While this information offers valuable insights in its abundance, it also underscores the challenges of large-scale analyses. For example, the use of less stringent aligners can lead to the generation of extensive collections of matches, making it more difficult to select the most similar link. LMEMs help address this challenge by selecting both maximum coverage and identity. As *C. hyointestinalis subsp. hyointestinalis* is an emerging pathogen^35^ identifying MGEs and their genomic links with Graphite can help in understanding its genomic evolution and potentially inform surveillance strategies.

While the genetic exchange between different *Campylobacter* species warrants further investigation, the power of Graphite to rapidly detect such events in this example dataset showcases its flexibility in identifying DNA transfer events. The examples above include previously undetected transfers between genomes and chromosomes, showing that LMEM analysis is a treasure trove for evolutionary genomics analysis, and underscoring the value of fast and memory-efficient algorithms to enable exploration of the vast microbial sequence space.

## Acknowledgments

This research was supported by ZonMw under project number 541003001, the European Research Council (ERC) Consolidator grant 865694: DiversiPHI, the Deutsche Forschungsgemeinschaft (DFG, German Research Foundation) under Germany’s Excellence Strategy – EXC 2051 – Project-ID 390713860, and the Alexander von Humboldt Foundation in the context of an Alexander von Humboldt-Professorship founded by German Federal Ministry of Education and Research. Special thanks to the members of the Theoretical Biology and Bioinformatics group at Utrecht University and the Viral Ecology & Omics Group at Friedrich Schiller University Jena, as well as to Yasas Wijesekara from the University of Medicine, Greifswald, for valuable input to develop the algorithm.

## Online Methods

We describe the Graphite algorithm which finds the longest maximum exact matches (LMEMs) between one or multiple queries and a reference database. Graphite builds on compacted colored De Bruijn graphs (ccDBGs)^48^ and suffix arrays (SAs)^49^. These data structures are briefly described in the Supplementary Information. The ccDBG input format is a reduced GFA to save storage space, which we here produced with Cuttlefish (REF). Alternatively, the input can be derived from a regular GFA file. After Graphite takes a ccDBG and a set of query identifiers, the algorithm consists of three phases: query SA construction, reference alignment, and Graphite output generation.

### Query suffix array construction

Before efficiently searching the ccDBG for LMEMs we have to convert it into an SA. Similar to bf-MEM, we construct the SA from queries instead of references to avoid indexing portions of the graph that may not be used. To index the SA using the SA-IS algorithm^50^, node identifiers for all query paths should be concatenated. For textual data, the “$” character is often used to separate different queries as it is lexicographically smaller than the alphabet letters and therefore does not affect sorting. For numerical data, we used negative numbers each identifying the query following it. In regular ccDBGs, node identifiers can also be negative depending on the forward/reverse direction that a sequence traverses through it, which may conflict with negative separators. To resolve this and facilitate reverse complement matching, we use the second-to-last bit of the node identifier to indicate the traversal direction (**Figure 5**). This caps the node identifiers at 2^30^ instead of 2^31^ which is validated when reading the graph file. After constructing the SA, the longest common prefix (LCP) and inverse SA (ISA) can be derived using standard algorithms^22^. The steps to construct the data structures from a graph file are given in **Algorithm 1**.

#### Algorithm 1. Suffix array construction from graph file

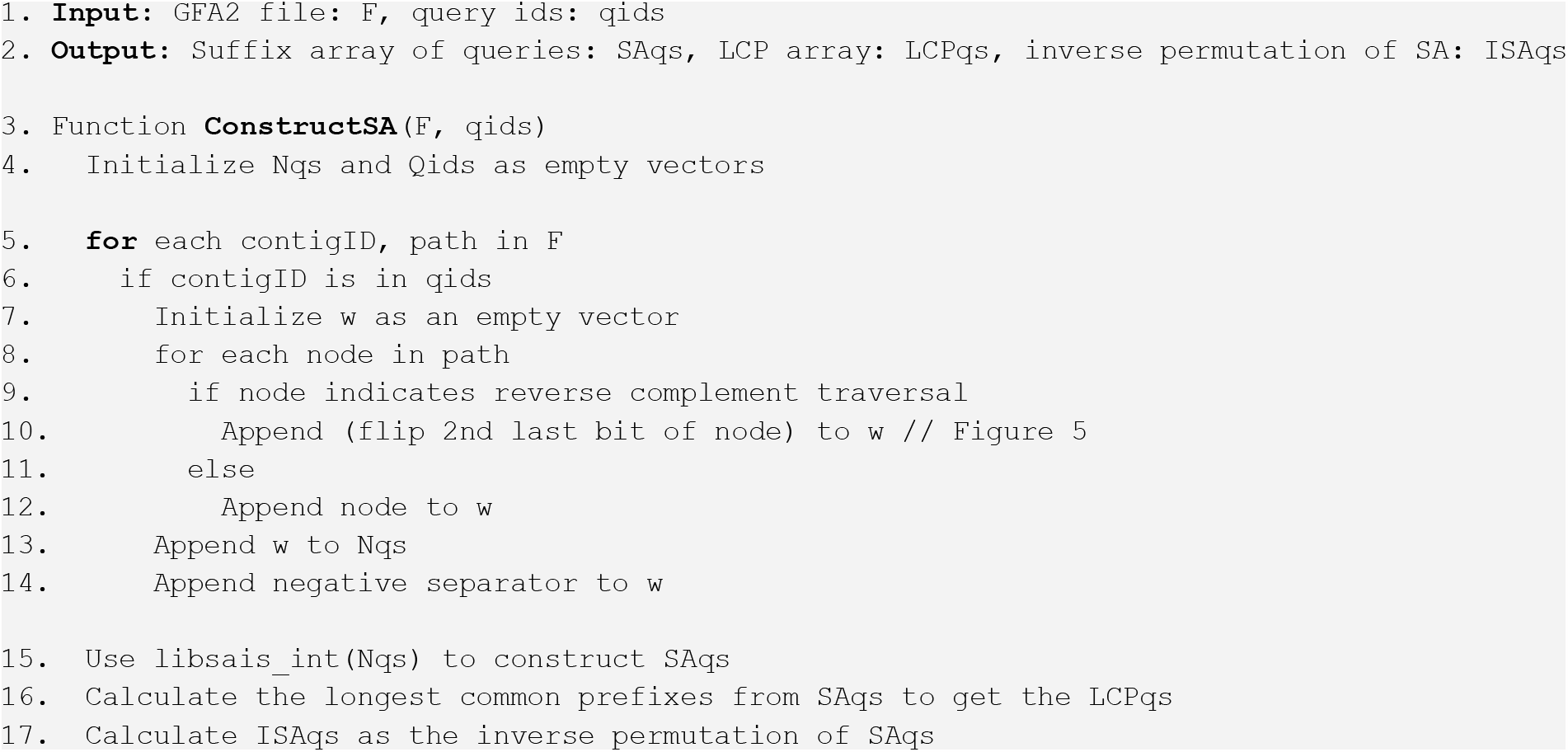

### Reference alignment

After constructing the SA of queries, we efficiently match reference sequence suffixes to identify LMEMs. Graphite processes reference nodes similarly to queries, converting reverse complement nodes to a positive number by using the second last bit. We locate MEMs between references and queries using the ISA and LCP (**Algorithm 2**). To find reverse complement matches, we reverse the reference path, flip the second last bit of the nodes, and then apply **Algorithm 2** again. For each identified MEM, we compare it to previously found LMEMs and replace them if the new MEM is longer (**Figure 6**). Since nodes in the compacted graph can represent sequences of varying lengths, we calculate MEM length by summing node lengths and removing their k-1 overlap. After iterating over all references, we obtain all LMEMs between queries and references.

#### Algorithm 2. Aligning references to the SA to discover LMEMs

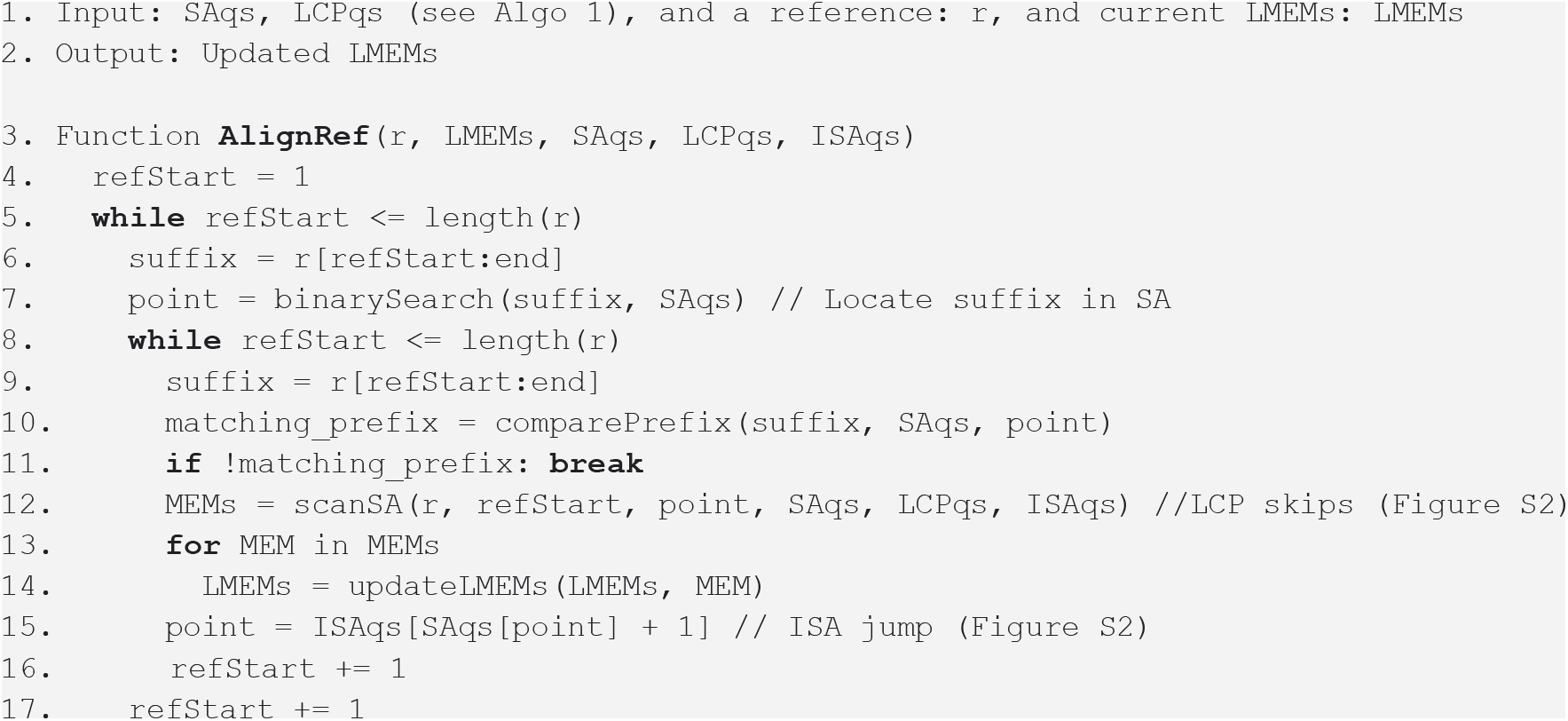

### Graphite output generation

We generate a tab-separated file containing the query identifier, the reference identifier, the LMEM start position, the LMEM end position, and the original MEM size. As the original MEM might be trimmed, its size compared to the LMEM size provides insight into the extent of trimming (**Figure 6**).

### Updating LMEMs and speed-up algorithm

In our algorithm description, we omitted details on optimizing LMEM updates. LMEM sizes for each query node are stored in LMEMsizes (**Algorithm 3**, line 16), while their associated reference identifiers are in LMEMorigins (line 17). For example, an LMEM with reference X for nodes [A, B, C] of 35 nucleotides (nts) each is stored as LMEMsizes = [35, 35, 35] and LMEMorigins = [X, X, X]. This enables efficient updates of the flank overlaps. When a new MEM, 50 nts, with reference Y overlaps with the last node C, at position j of the previous LMEM, we observe that LMEMsizes[j] < length(MEM). Consequently, we update part of the LMEM, resulting in LMEMsizes = [35, 35, 50, 50, 50] and LMEMorigins = [X, X, Y, Y, Y]. However, for long matches, comparing many positions for each MEM becomes computationally expensive, especially when searching all suffixes. To address this we apply **Algorithm 3** where we only update LMEMs after checking for a longer or equally long LMEM.

#### Algorithm 3. Updating LMEMs

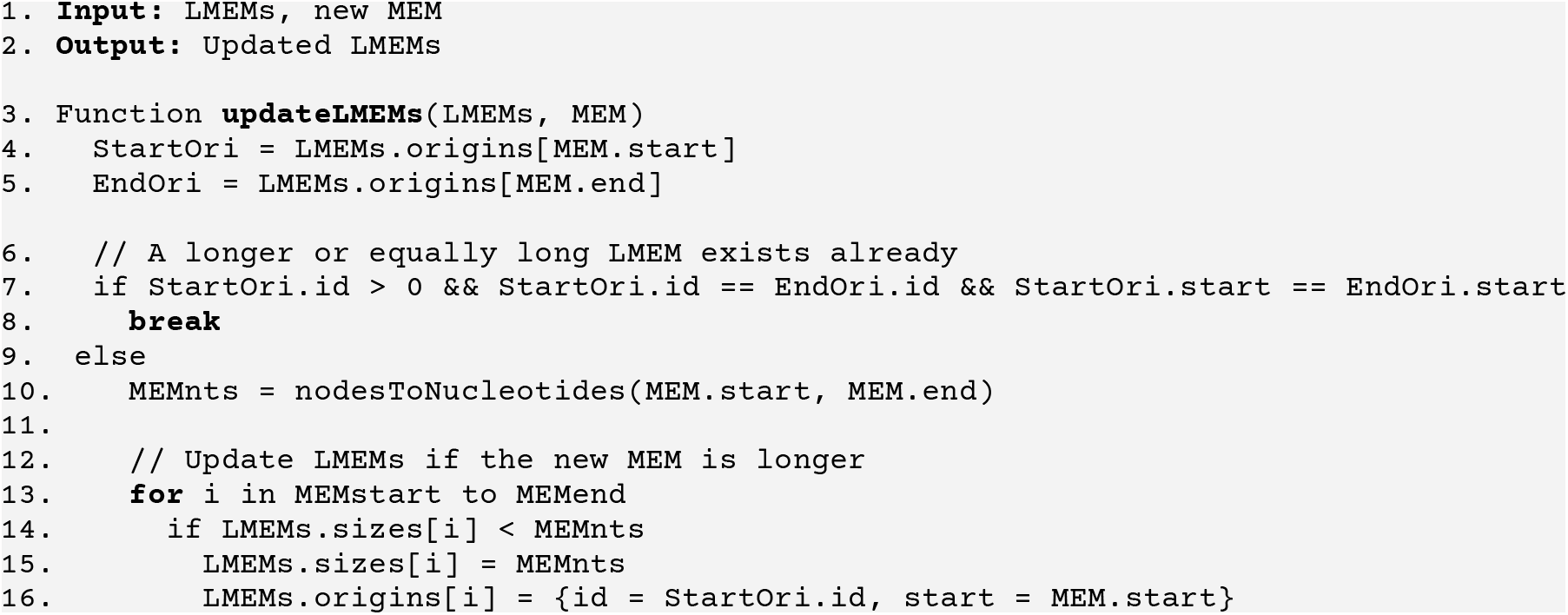

### Graphite validation and application to existing datasets

We collected all complete *Campylobacter* genomes, excluding those shorter than 1.5MB, from BV-BRC^36^ and built a MashTree^51^ that was rooted on the ancestor of *C. coli* and *C. jejuni* and visualized with iTOL^52^. We re-annotated species when their BV-BRC annotation deviated from that of the majority in the monophyletic clade (see **Table S1**). To use Graphite, we built a ccDBG of all *Campylobacter* genomes with Cuttlefish ^33^ (k-mer size: 31) and saved storage space using reduced GFA (-f 3) output format.

LMEM selection by Graphite was validated on three *C. jejuni* genomes (CP071576.1, CP071584.1, CP071578.1) and compared to E-MEM results with match size 31 (-l -b). Correct LMEMs were determined by their maximum length among all MEMs covering a position (**Figure 1**).

To evaluate Graphite’s performance, we compared it to E-MEM^25^ and bf-MEM^26^. We queried *C. coli* genomes against all other *Campylobacter* genomes using all three tools. E-MEM used the same settings as earlier, while bf-MEM used a minimum match size of 33 (-l) and a k-mer size of 32 (-k). This choice was due to restrictions in bf-MEM, requiring k%4 == 0 and l > k. The computer setup was consistent: Gold-6240R CPU, 500GB RAM, and HDD storage drive.

To demonstrate Graphite’s ability to efficiently detect DNA exchange, we applied it to all *Campylobacter* genomes, comparing each genome to the others, and highlighting striking examples of HGT. Directly visualizing the LMEM region can result in an explosion of paths and nodes obfuscating the actual shared region and context. To simplify, in **Figure 4** we visualized LMEMs using parameters for example n = 0.9 and l = 1000, where N determines the minimum fraction of nodes overlapping the LMEM for inclusion, and L specifies the number of extending nodes on each side. See https://github.com/rickbeeloo/subgraphs for the implementation. The simplified subgraph was then visualized using Bandage^53^ and gene plots were created using DNA Features viewer^54^

## Notes

https://github.com/MGXlab/Graphite

### Competing Interest Statement

The authors have declared no competing interest.

