## Supplementary data/figures for "Graphite: painting genomes using a colored De Bruijn graph"

### Supplementary Information

#### Compacted colored De Bruijn graphs

A sequence collection can be losslessly compressed using a compacted colored De Bruijn graph (ccDBG, **Figure S1**). Each genome follows a path through the nodes in the ccDBG that spell out its sequence. As nodes represent both the forward and reverse complement sequences (**Figure S1B**) a negative sign after the node identifier is used to indicate a reverse complement traversal (**Figure S1C**). When finding maximum exact matches (MEMs) between two sequences, these can occur in the same orientation or reverse complement. Forward MEMs show an identical stretch of nodes between two sequences, for example [2+, 3+] as shown in **Figure S1C**. This can also include mixed signs like [1+, 2-, 3+]. Reverse complement MEMs should share the same node identifiers but with one path being in reverse order with all signs flipped. An example is shown in **Figure S1D**, where [1+, 2+] matches [2-, 1-], and [2-, 3+] matches [3-, 2+]. The negative signs and matching rules become important later for the construction of suffix arrays and identifications of the LMEMs.

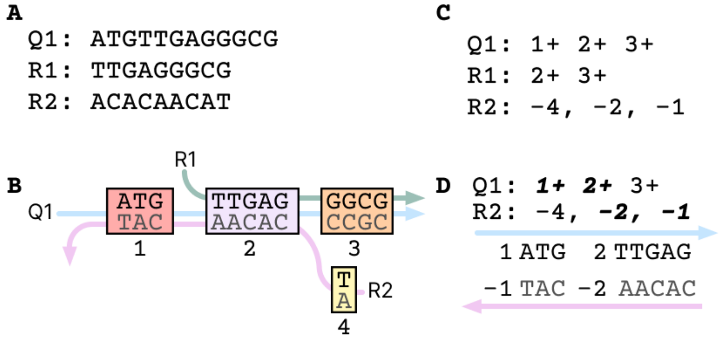


***Figure S1. Colored De Bruijn graph illustration.*** Panel **A** shows three hypothetical sequences and **B** shows a ccDBG representing them. Each node has a unique identifier that represents a sequence and each genome follows a path through these nodes that spell out the genome (**C**). For simplicity, we did not show the k-1 overlap between nodes. A feature of the ccDBG is that a node represents both the forward and reverse complement sequences. This allows for compression as only the forward sequence needs to be saved (black sequences in **B**) and a sign is used to indicate the reverse complement (gray sequences in **B**). An additional advantage of this way of storing graphs is that reverse complement MEMs are directly evident from shared node identifiers even when the raw sequences are different (**D**).

#### Suffix array and derived data structures

A suffix array (SA) is an array of all the suffixes of a given string in lexicographical order (**Figure S2**). To reduce space, it does not hold the actual suffixes, but just the start position of the suffix in the original string. The lexical ordering allows us to place any query string in the SA using a binary search (**Figure S2A,** right). For example, when searching ABA in the sequence ABABCD the binary search will stop at the second suffix, ABABCD (**Figure S2A**, red arrow).

Since they are sorted, the suffixes that share a prefix are grouped together and we can move up or down from the binary search point to locate other matches. To skip unnecessary character comparisons while searching, we store the LCP between subsequent suffixes (**Figure S2B**). To locate the second suffix of our query ABA, namely BA, we can leverage the ISA (**Figure S2C**) to directly jump to the match locations in the SA, namely BABCD. From here we can again scan up and down using the LCP to find all other matches with BA. In this case, a one-character match for B with the suffix BCD above it. This shows that we can use the suffix array, LCP, and ISA to efficiently locate a sequence and all its matches.

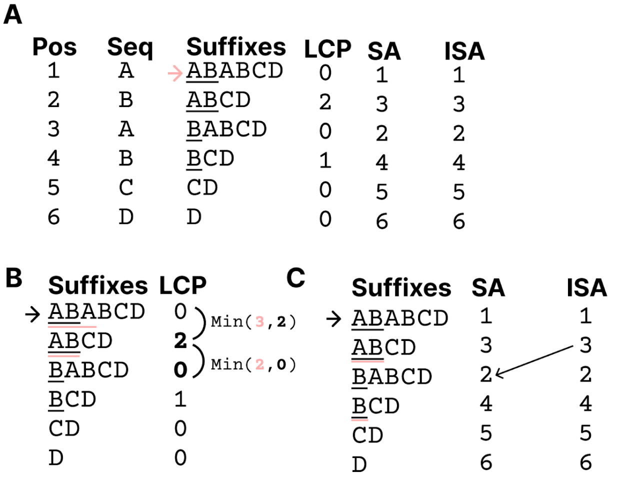


***Figure S2. Examples of a suffix array, inverse suffix array, and longest common prefix array.*** Given a sequence (Seq) ABABCD, we can sort all its suffixes lexicographically (**A**) and store the starting position of each suffix in the original string to obtain the suffix array (SA). The sorted nature of the SA allows us to use a binary search to place a string, such as "ABA", in the SA (red arrow in **A**). The longest common prefix (LCP) array describes the length of the shared prefix between two consecutive suffixes. For example, the suffix at position 2 shares "AB" with the suffix at position 1, and hence has an LCP of two (**B**, black underlined letters). Given a match at any location in the SA, we can thus directly infer matches with flanking suffixes by leveraging the LCP as we know that we can always match at least $Min(match_{size}, LCP)$ characters. Continuing the example of "ABA" after its match with the first suffix we can calculate its match with the second suffix by using the LCP (**B**, red underlined letters). Since our query matches the first three characters of the first suffix, we know that if we search for the next suffix in the query, "BA" it could also match two characters with the next suffix of our match "ABABCD", namely "BABCD". To locate "BABCD" in the SA we can follow the ISA according to $ISA[SA[index] + 1]$. Filling that in for the first suffix in the SA gives us $ISA[SA[1] + 1] = ISA[1 + 1] = ISA[2] = 3$ and indeed $SA[3]$ = "BABCD" (**C**).

**Figure S3. MashTree of campylobacter genomes.** A MashTree of 576 *Campylobacter* genomes colored according to the BV-BRC species annotation. Manually corrected species are shown with red arrows. See **Supplementary Table S1** for all details on all strains.

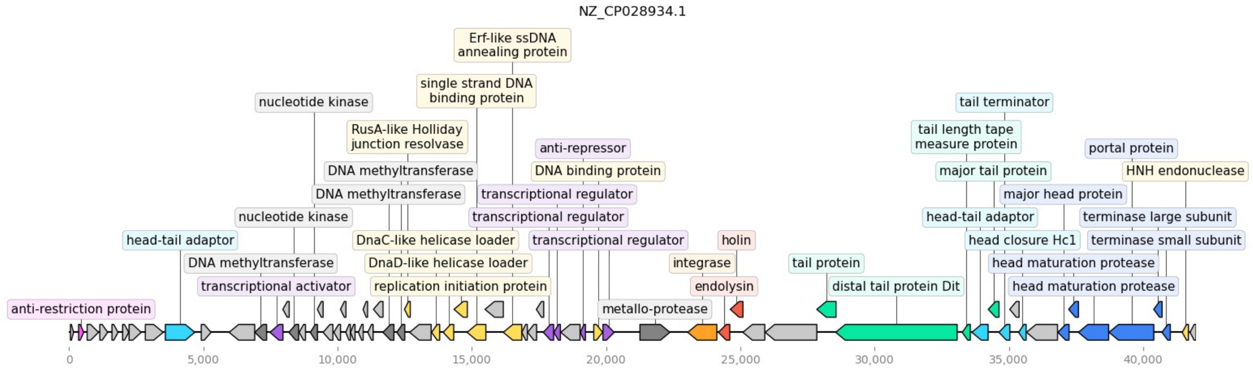


**Figure S4 Gene annotation of CP028934.1.** CP028934.1 is annotated as a plasmid but when annotated with HMMs from PHROGs [55] we found typical bacteriophage genes including structural and replication clusters, integrase, and a terminase large subunit.

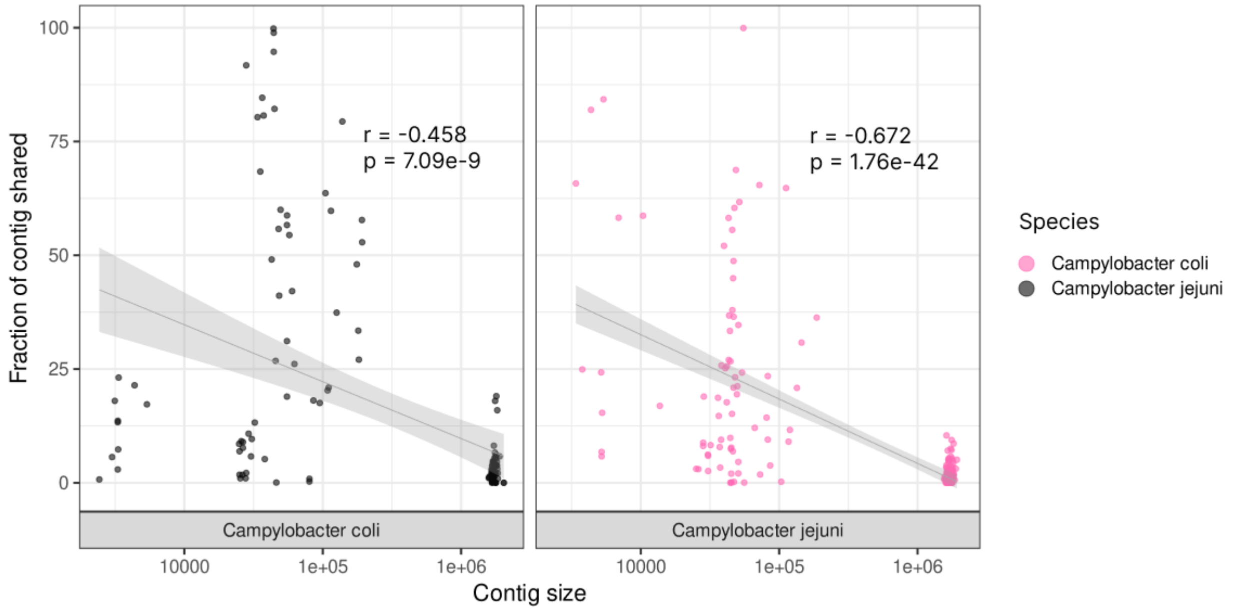


***Figure S5. Percentage of contig shared between C. coli and C. jejuni.*** We selected all LMEMs shared between *C. coli* and *C. jejuni* and calculated the percentage of each contig covered by LMEMs of the other species. Generally, shorter contigs (≤100,000) shared a higher fraction of the contig with the other species. For the longer contigs, around the length of chromosomes (>1,500,000), up to 19% of the sequence of *C. coli* contigs could be shared with *C. jejuni*. The Pearson correlation (r) and its p-value are shown.
